# Evaluation of flow cytometry for cell count and detection of bacteria in biological fluids

**DOI:** 10.1101/2021.10.04.463144

**Authors:** N. Dossou, I. Gaubert, C. Moriceau, E. Cornet, S. Le Hello, D. Malandain

## Abstract

The analysis of biological fluids is crucial for the diagnosis and monitoring of diseases causing effusions and helps in the diagnosis of infectious diseases. The gold standard method for cell count in biological fluids is the manual method using counting chambers. The microbiological routine procedures consist of Direct Gram staining and culture on solid or liquid media. We evaluate the analytical performance of SYSMEX UF4000 (Sysmex, Kobe, Japan) and Sysmex XN10 (Sysmex, Kobe, Japan) in comparison with cytological and microbiological routine procedures. A total of 526 biological fluid samples were included in this study (42 ascitic, 31 pleural, 31 peritoneal, 125 cerebrospinal, 281 synovial, and 16 peritoneal dialysis fluids). All samples were analysed by flow cytometry and subsequently processed following cytological and/or microbiological routine procedures. With regards to cell counts, UF4000 (Sysmex, Kobe, Japan) showed a performance which was at least equivalent to those of the reference methods and superior to those of XN10 (Sysmex, Kobe, Japan). Moreover, the bacterial count obtained with UF4000 (Sysmex, Kobe, Japan) was significantly higher among culture or Direct Gram stain positive samples. We established 3 optimal cut-off points to predict Direct Gram stain positive samples for peritoneal (465.0 bacteria/μL), synovial (1200.0 bacteria/μL), and cerebrospinal fluids (17.2 bacteria/μL) with maximum sensitivity and negative predictive values. Cell count and detection of bacteria by flow cytometry could be used upstream cytological and microbiological routine procedures to improve and accelerate the diagnosis of infection of biological fluid samples.

## Introduction

The analysis of biological fluids is crucial for the diagnosis and monitoring of diseases causing effusions and helps in the diagnosis of infectious diseases. The gold standard method for cell count in biological fluids is the manual method using counting chambers^1^. The current REMIC recommendations for microbiological procedures consist of Direct Gram staining (DGS), culture on solid media, liquid media, and/or incubation in blood culture bottles. These manual methods require significant expertise and technical time.

Cytological and microbiological analysis of effusion fluids, cerebrospinal fluids, and synovial fluids should be performed promptly because of the rapid degradation of cells, especially neutrophils. In addition, the results of these analyses are important tools for clinical decision-making, particularly in the case of infectious diseases, where the early initiation of appropriate anti-infectious treatment results in a better prognosis^2,3,4^.

The automation of these analyses has several advantages over manual methods, due to its speed and the lack of a need for sample pre-treatment. Being operable 24/7 and offering reliable analysis, the SYSMEX XN10 (Sysmex, Kobe, Japan) and the SYSMEX UF4000 (Sysmex, Kobe, Japan) machines could be an interesting alternative to manual methods.

The aim of this study was to evaluate the analytical performance of the SYSMEX UF4000 (Sysmex, Kobe, Japan) and SYSMEX XN10 (Sysmex, Kobe, Japan) as methods for the cytological analysis of different body fluids in comparison to manual methods: cell count in KOVA counting chambers and differential leukocyte count after staining with May Grunwald Giemsa (MGG).

The secondary aim of this study was to evaluate the analytical performance of the SYSMEX UF4000 (Sysmex, Kobe, Japan) as a method for the microbiological analysis in comparison with DGS and/or the conventional culture on solid media, liquid media and enrichment broth.

## Materials and methods

### 1/ Sample inclusion

In total, 538 biological fluid samples received at the Microbiology or Haematology Departments of Caen University Hospital between June 2020 and July 2021 were included in the study. The samples were collected in sterile tubes with no chemical preservatives and processed immediately after arrival.

#### Comparison of the performance of cytological analysis methods

Six different kinds of biological fluids were processed by the routine cytological procedures and subsequently by the SYSMEX XN (Sysmex, Kobe, Japan) and the SYSMEX UF4000 (Sysmex, Kobe, Japan): ascitic fluid (AF), pleural fluid (PF), peritoneal fluid (PRF), peritoneal dialysis fluid (PDF), synovial fluid (SF) and cerebrospinal fluid shunt (CSF). The SF was previously diluted in saline solution before analysis (1:10 dilution). Specimens with insufficient volume (less than 1 mL) were excluded (data not recorded).

Some clinical and biological information was sought in the patient files: medical reason for the punction of biological fluids and reported history of haematological malignancy.

#### Comparison of the performance of microbiological analysis methods

Six different kinds of biological fluids were processed by the routine microbiological procedures and subsequently by the SYSMEX UF4000 (Sysmex, Kobe, Japan): AF, PF, PRF, PDF, SF and CSF. The SF was previously diluted in saline solution before analysis (1:10 dilution). Specimens with insufficient volume (less than 1 mL) were excluded (data not recorded).

Some clinical and biological information was sought in the patient files: clinical symptoms, anti-infectious treatment and diagnosis of infection.

### 2/ Standard methods

#### Routine cytological procedures

Cell counting (red blood cells and leukocytes) of body fluids was done manually, using ten cell count slides with grids (KOVA® Glasstic® Slide 10 [CML, Nemours, France]). The cells were quantified under x10 and x40 magnifications. The results were reported as the number of cells per microliter.

The differential leukocyte count of the body fluids was done manually after making a spot with cytospin and then staining with MGG. The differential count was performed at x100 magnification on a total count of 100 cells with the differentiation of polynuclear cells and mononuclear cells.

#### Routine microbiological procedures

A DGS was performed in centrifugation pellets (10 min at 3000 rpm) for all PRF, PF, SF, and, in bloody and/or mucous PDF. For non-mucous LDP, the DGS was performed after making a cytospin spot (2 drops, 5 min). A DGS on the primary sample was performed for all AF and CSF with sufficient volume and with leukocyte count greater than or equal to 10/mm^3^.

Biological fluids were inoculated manually. PDF and AF were added into aerobic and anaerobic blood cultures bottles. These bottles were incubated into the BACT/ALERT® VIRTUO® (BioMérieux, Marcy-L’étoile, France) for 7 days. CSF were inoculated on chocolate agar (BioMérieux, Marcy-L’étoile, France) and on Brain Heart Infusion medium (BioMérieux, Marcy-L’étoile, France). PF were inoculated on blood agar (BioMérieux, Marcy-L’étoile, France), chocolate agar (BioMérieux, Marcy-L’étoile, France) and on Schaedler medium (BioMérieux, Marcy-L’étoile, France). PRF were inoculated on blood agar (BioMérieux, Marcy-L’étoile, France), chocolate agar (BioMérieux, Marcy-L’étoile, France), Bromocresol purple agar (BioMérieux, Marcy-L’étoile, France), Schaedler medium (BioMérieux, Marcy-L’étoile, France), and on selective chocolate agar TM+PolyViteX VCAT3 (BioMérieux, Marcy-L’étoile, France) for female patient. SF were inoculated on chocolate agar (BioMérieux, Marcy-L’étoile, France), blood agar (BioMérieux, Marcy-L’étoile, France) and incubated on blood cultures bottles into the BACT/ALERT® VIRTUO® (BioMérieux, Marcy-L’étoile, France) for 14 days. Bacterial identification was achieved using MALDI-TOF MS (Matrix associated laser desorption/ionization time-of-flight mass spectrometry) (Bruker, Bremen, Germany).

### 3/ Automated methods compared

#### Sysmex UF4000 (Sysmex, Kobe, Japan)

SYSMEX UF4000 (Sysmex, Kobe, Japan) is an automated urine and body fluid analyser whose technology is based on flow fluorocytometry with a blue laser. It provides 9 parameters for body fluids: erythrocyte count (RBC), total nucleated cell count (TNC), epithelial cell count, white blood cell count (WBC), count and percentage of mononuclear cells (MN#, MN%), count and percentage of polynucleated cells (PN#, PN%), and the count and classification of bacteria (BACT). Counting and classification is performed in three steps: cell labeling (nucleic acids, membrane components or surface proteins), hydrodynamic focusing and finally characterisation by the blue laser using four different signals. The FSC, SFL, SSC and DSS signals, respectively, provide information for particle size, fluorescence, internal complexity and depolarisation.

All samples were analysed by flow cytometry using the Sysmex UF4000 (Sysmex, Kobe, Japan) and following the manufacturer’s recommendations. Samples were analysed manually using the STAT mode and the required minimum sample volume was 0.6mL. High and low positive controls were processed twice daily.

#### Sysmex XN (Sysmex, Kobe, Japan)

Sysmex XN (Sysmex, Kobe, Japan) is an automated blood samples analyser. It is made of different modules. The classic module performs blood counts. The expert modules perform fluorescence counts of blood platelets and reticulocytes. The “Body fluid” module performs blood fluid samples analysis by using two channels. The RET channel performs red blood cell counts by impedance. The WDF channel performs white blood cell counts by flow fluorocytometry. It provides different parameters: total nucleated cell (TC-BF), white blood cell count (WBC-BF), percentage of mononuclear cells, and percentage of polynucleated cells.

Samples were analysed on the SYSMEX XN (Sysmex, Kobe, Japan) after their analysis on the SYSMEX UF4000 (Sysmex, Kobe, Japan) following manufacturer’s recommendations. Samples were analysed manually on the “body fluid” module. The required minimum sample volume was 0.3mL. Controls were processed twice daily.

### 4/ Statistical analysis

Statistical analyses was performed using EXCEL^®^ and XL STAT^®^.

#### Comparison of the performance of cytological analysis methods

Red and white blood cell counts were compared using the Spearman’s correlation coefficient and the non-parametric Wilcoxon test for the comparison of two dependent variables.

Differential leukocyte counts were compared using the Spearman’s correlation coefficient and the Rümke table. The differences were considered statistically significant with a *p*-value < 0.05.

#### Comparison of the performance of microbiological analysis methods

The different parameters measured by flow cytometry (FCM) were defined as dependent variables using the culture and DGS results as independent variables. The distribution of independent variables across groups was compared using the Mann-Whitney U test for the comparison of two independent variables. The differences were considered statistically significant with a p-value < 0.05.

The receiver operating characteristic (ROC) curves were used to compare the UF4000 bacterial counts to the culture and DGS results obtained by routine procedures. The cut-off values were chosen based on best balance between sensitivity and specificity, giving priority to sensitivity in order to detect DGS-positive samples and giving priority to specificity in order to detect culture positive samples.

### 5/ Ethical statement

The study was carried out without any additional intervention in patients. All samples were processed following routine cytological and microbiological procedures. Patient data were anonymised before analysis.

Demographic, clinical, and biological data were obtained from the laboratory information system (TD NEXLABS^©^) and the hospital information system (REFERENCE^©^).

## Results

### 1/ Comparison of the performance of cytological analysis methods

A total of 189 biological fluids samples were analysed by FCM on the SYSMEX UF4000 (Sysmex, Kobe, Japan) and the SYSMEX XN (Sysmex, Kobe, Japan). They were subsequently processed by the routine cytological procedures (white blood cell and red blood cell count in KOVA counting chambers, manual differential leukocyte count after staining with MGG). Seven samples were excluded due to a lack of clinical or biological information. Finally, 182 samples originating from 151 patients were included in this part of the study: 42 AF, 31 PF, 31 PRF, 16 PDF, 30 SF and 32 CSF. The mean age was 60 years (Interquartile range (IQR): 22) and the sex ratio was 1.43. The characteristics of the included patients are shown in **Table 1.**

**Table 1.** Characteristics of included patients for comparison of the performance of cytological analysis methods.

|  | All fluids | AF<sup>a</sup> | PF<sup>b</sup> | PRF<sup>c</sup> | PDF<sup>d</sup> | SF<sup>e</sup> | CSF<sup>f</sup> |
| --- | --- | --- | --- | --- | --- | --- | --- |
| **Total** | 182 | 42 | 31 | 31 | 16 | 30 | 32 |
| **Age**<br>(mean ± standard deviation) | 60.47 ± 19.91 | 64.76 ± 15.49 | 63.84 ± 17.34 | 49.10 ± 24.78 | 51.56 ± 29.23 | 69.17 ± 12.80 | 58.91 ± 16.18 |
| **Sex ratio** | 1.43 | 2.23 | 1.07 | 1.38 | 1.29 | 2.33 | 0.78 |
| **Hematologic malignancy, n, (%)** | 13 (7.14%) | 7 (16.67%) | 5 (16.13%) | 1 (3.23%) | 0 (0.00%) | 0 (0.00%) | 0 (0.00%) |
| **Antimicrobial treatment, n, (%)** | 59 (32.42%) | 11 (26.19%) | 6 (19.35%) | 14 (45.16%) | 8 (50.00%) | 5 (16.67%) | 15 (45.88%) |

<sup>a</sup> AF: Ascitic fluid, <sup>b</sup> PF: Pleural fluid, <sup>c</sup> PRF: Peritoneal fluid, <sup>d</sup> PDF: Peritoneal dialysis fluid, <sup>e</sup> SF: Synovial fluid, <sup>f</sup> CSF: Cerebrospinal fluid shunt.

#### a) Cell count

The three methods compared in this study have different limits of quantification for red blood cell (RBC) and white blood cell (WBC) counts; they are shown in **Table 2**. Red blood cell counts obtained by FCM using the SYSMEX XN (Sysmex, Kobe, Japan) were rounded to 1000/μL.

**Table 2.** Limits of quantification for red blood cell and white blood cell counts obtained by automated methods UF and XN, and by standard procedure.

|  | Sysmex XN10 | Sysmex UF4000 | KOVA |
| --- | --- | --- | --- |
| **Red blood cell (RBC)** | 0 – 10 × 10<sup>6</sup>/μL | 15 – 100 000/μL | 0 – 1000/μL |
| **White blood cell (WBC)** | 0 – 5000 × 10<sup>3</sup>/μL | 2 – 10 000/μL | 0 – 1000/μL |

Comparison results of the automated methods to the KOVA standard method for RBC counts are shown in **Table 3** and in **Graph 1**. The comparison was performed on 96 samples. Eighty-six samples were excluded from the comparative analysis because standard method results were outside the limits of quantification (> 1000 red blood cells/μL). They also had results greater than > 1000 RBC/μL with the XN method, but 3 had results < 1000 RBC/μL with the UF method: 332/μL, 807.9/μL and 929.7/μL, respectively. The first two were pleural fluids taken from patients with pleural effusions, with no reported history of haematological malignancy or antibiotic therapy. The last was a peritoneal fluid sample taken from a patient with peritonitis symptoms and who was on broad-spectrum antibiotherapy.

**Table 3.** Comparison of red blood cell counts obtained by XN and UF methods to the standard method.

|  | Sysmex XN | Sysmex UF |
| --- | --- | --- |
| Mean difference | 46 | -15.16 |
| Lower limit of agreement | -926 | -154.14 |
| Upper limit of agreement | 1017 | 123.82 |
| Spearman's correlation coefficient | 0.5426<br>($p < 0.00001^*$) | 0.9616<br>($p < 0.00001^*$) |
| Regression equation | $Y = 0.218x + 125.32$ | $Y = 1.0341x + 9.7765$ |
| Paired Wilcoxon test (bilateral) | $p = 0.00089^*$ | $p = 0.11088$ |

*\*p-value < 0.05*
*All statistical tests were carried out in comparison with standard method (Red blood cell counts in KOVA counting chambers).*

**Graph 1.**
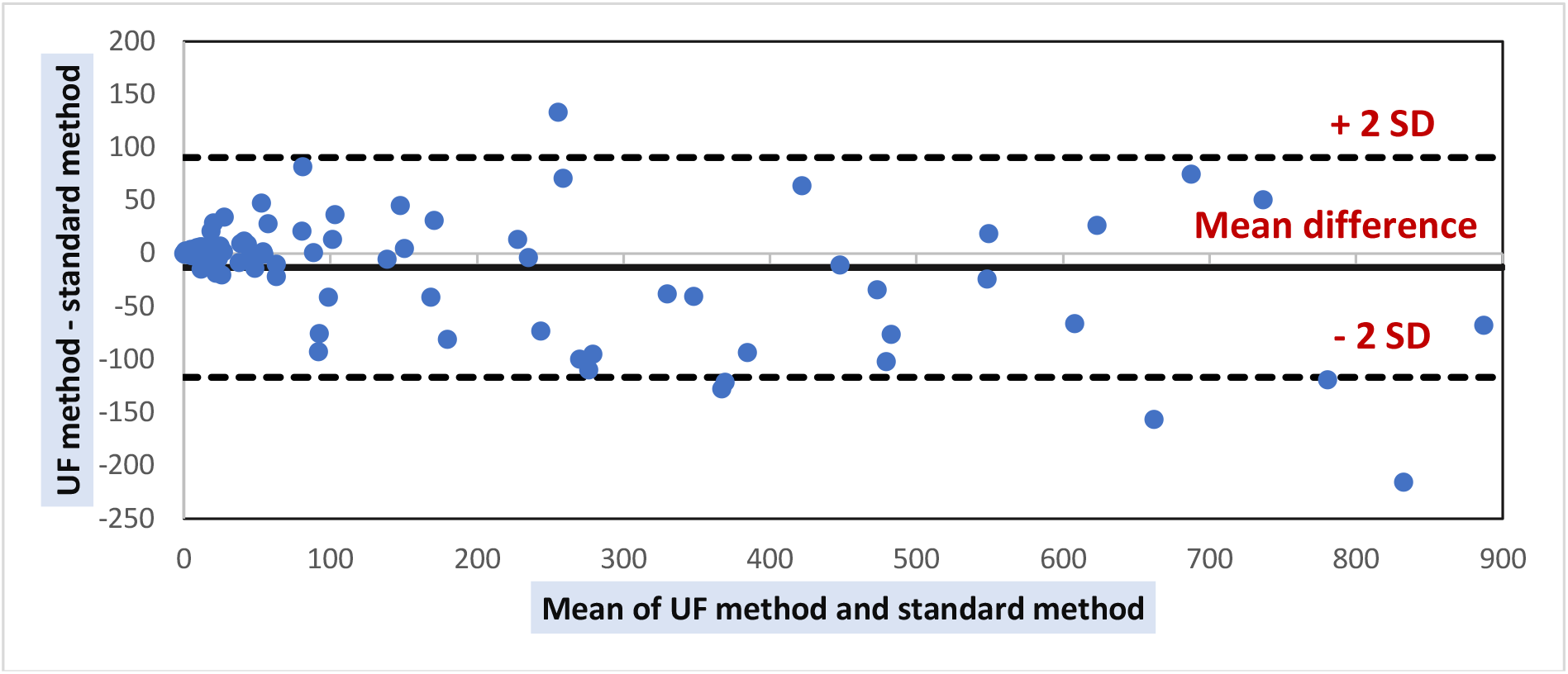
Bland Altman plot of differences between red blood cell counts by UF and standard method. *SD = standard deviation.

No significant correlation was found between XN and standard method results. The paired Wilcoxon test indicates that the results obtained by the two methods are significantly different. UF and standard method results are strongly correlated, and are not significantly different according to the paired Wilcoxon test.

Comparison of results of the automated methods and the KOVA standard method for WBC counts are shown in **Table 4, Graph 2** and in **Graph 3**. The comparison was performed on 135 samples. Forty-seven samples were excluded from the comparative analysis because standard method results were outside the limits of quantification (> 1000 WBC/μL). Two of them had results < 1000 WBC/μL with XN method (840/μL and 915/μL, respectively): one was peritoneal fluid from an organ donor and the other was peritoneal dialysis fluid. Also, two of them had results < 1000 WBC/μL with the UF method (995.7/μL and 454.9/μL, respectively): one was the same peritoneal fluid and the other was a peritoneal dialysis fluid taken from a patient who was on broad-spectrum antibiotherapy.

**Table 4.** Comparison of white blood cell counts obtained by XN and UF methods to the standard method.

|  | Sysmex XN | Sysmex UF |
| --- | --- | --- |
| Mean difference | 10.42 | 23.37 |
| Lower limit of agreement | -145.50 | -165.25 |
| Upper limit of agreement | 166.34 | 211.99 |
| Spearman's correlation coefficient | 0.955<br>($p < 0.00001^*$) | 0.9711<br>($p < 0.00001^*$) |
| Regression equation | $Y = 0.8829x + 8.9227$ | $Y = 0.9734x + 8.444$ |
| Paired Wilcoxon test (bilateral) | $p = 0.112932$ | $p = 0.0707247$ |

*\*p-value < 0.05*
*All statistical tests were carried out in comparison with standard method (Red blood cell counts in KOVA counting chambers).*

**Graph 2.**
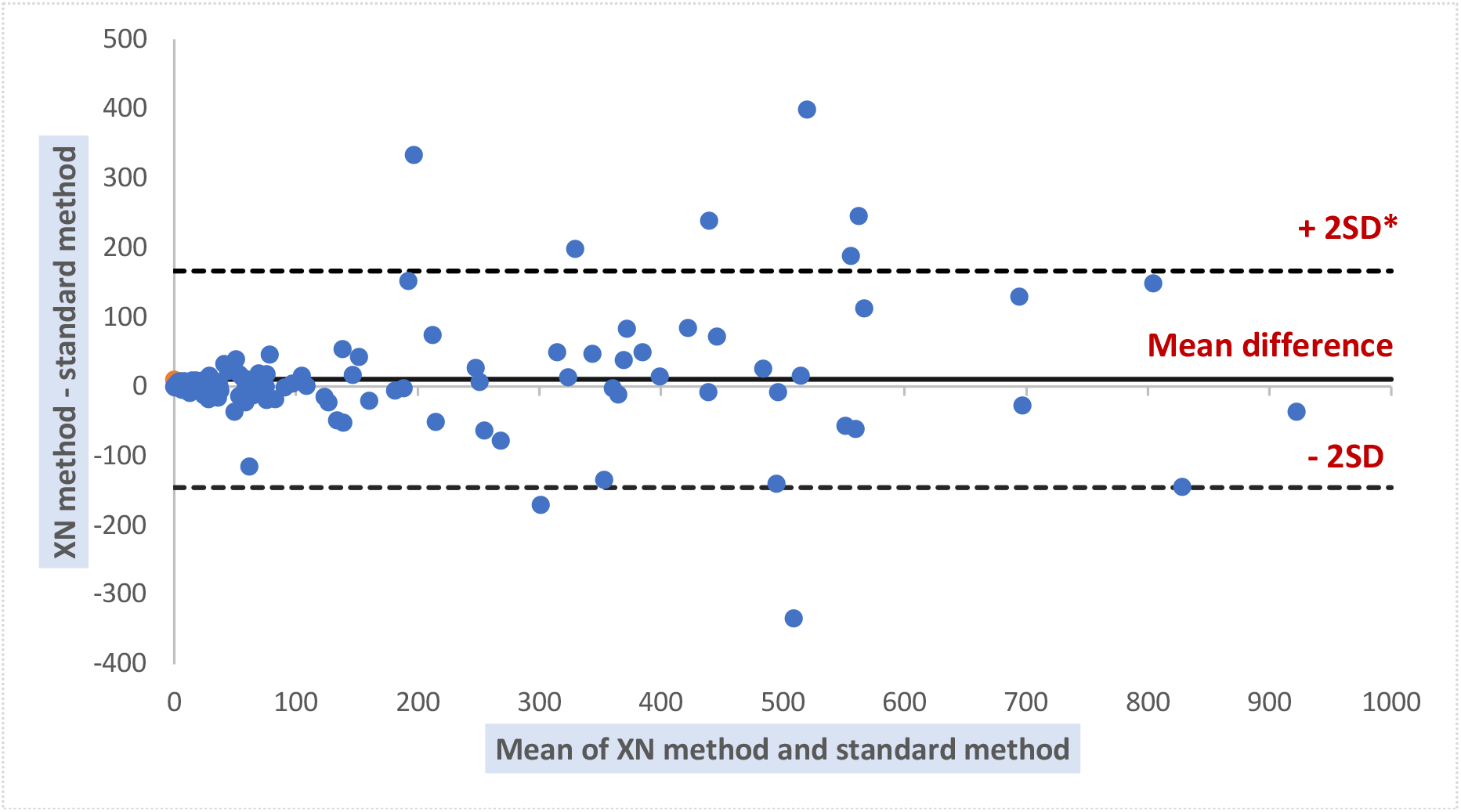
Bland Altman plot of differences between white blood cell counts by XN and standard method. *SD = standard deviation.

**Graph 3.**
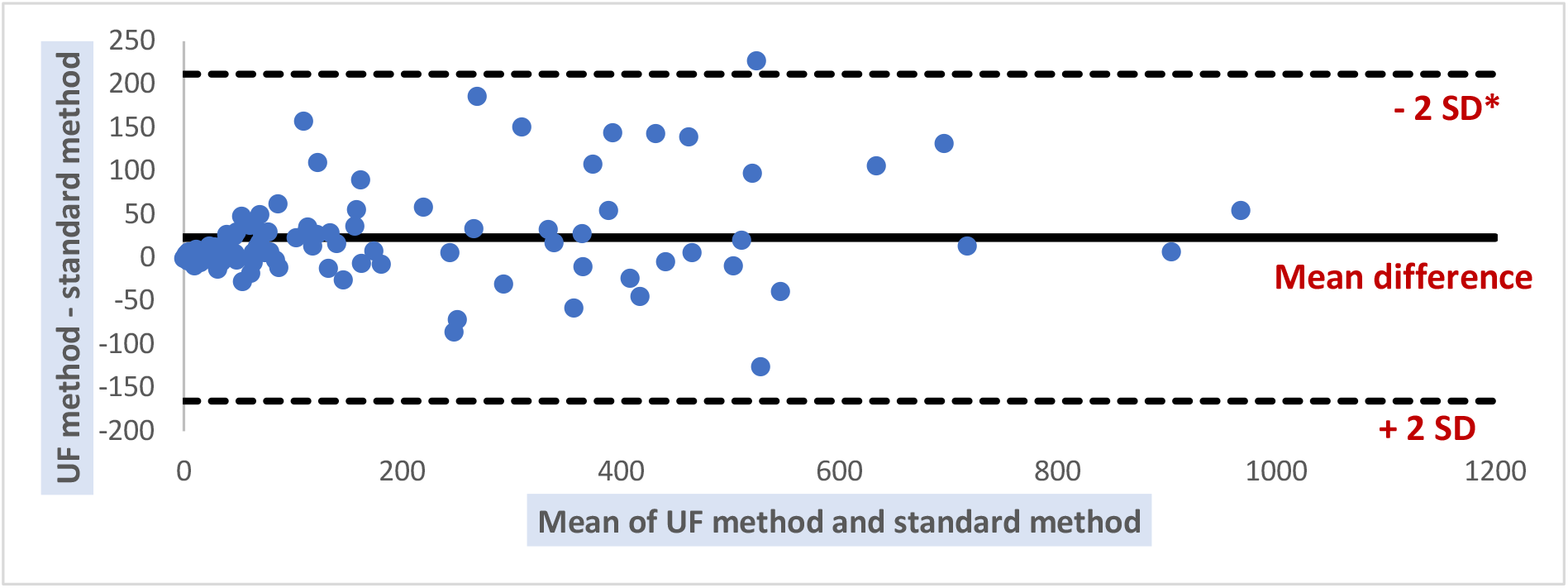
Bland Altman plot of differences between white blood cell counts by UF and standard method. *SD = standard deviation.

Strong and significant correlation was found between XN, UF and standard method results. The paired Wilcoxon test indicates that the results obtained by the different methods are not significantly different.

#### a) Differential leukocyte count

Comparison of the automated methods to the standard method for differential leukocyte counts was performed on 161 biological fluid samples. Twenty-one samples were excluded from the comparative analysis due to an insufficient amount of cells on the smear. Results show that the percentage of mononuclear cells obtained by the XN method (r^2^ = 0.9027, p-value < 0.0001) and by the UF method (r^2^ = 0.91, p-value < 0.0001) are strongly and significantly correlated with standard method results (**Graph 4**).

**Graph 4.**
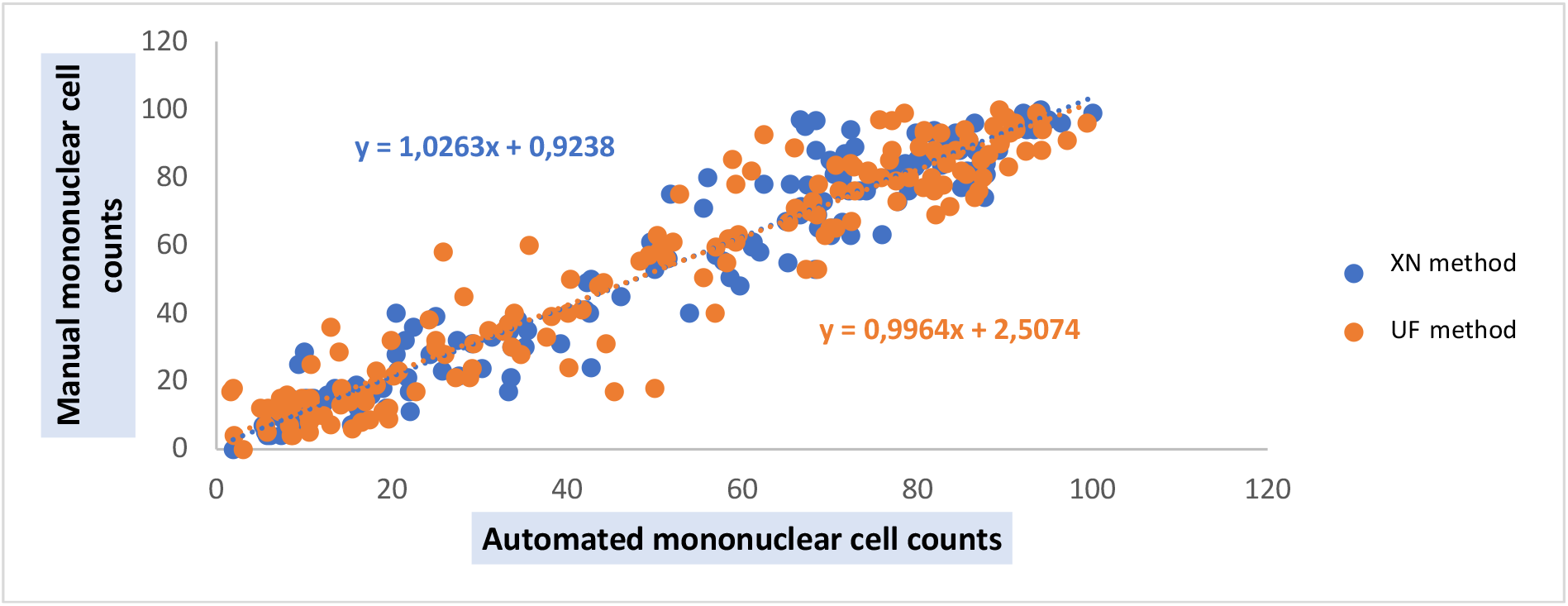
Regression lines between automated (UF et XN) and manual mononuclear cell counts.

In order to take into account the relative variations in leukocyte populations, we used the Rümke table to compare automated and manual mononuclear cell counts. Overall, 78.32% of the values obtained with the XN method lead to acceptable coefficients of variation in comparison with standard method results (**Graph 5**).

**Graph 5.**
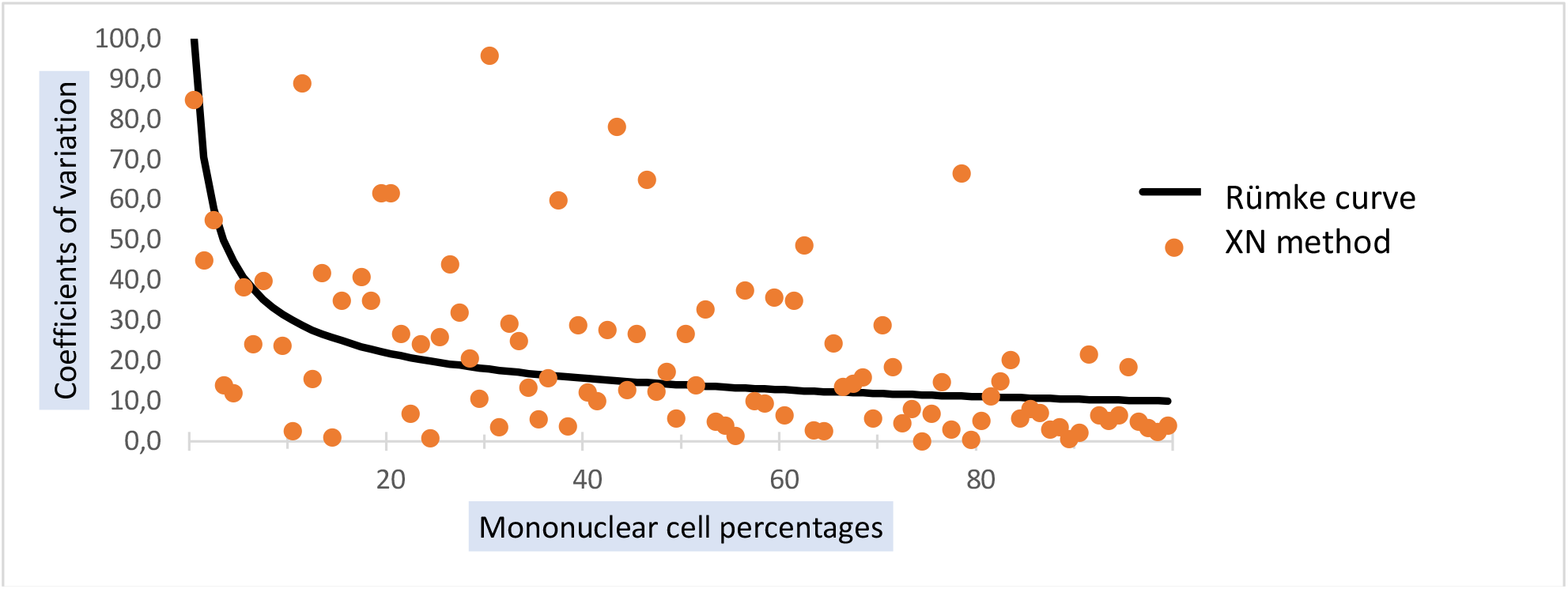
Comparison of mononuclear cell counts obtained by XN and standard methods. The Rümke curve was drawn using the Rümke table; it represents the limit value of acceptable coefficients of variation.

In addition, 84.25% of the values obtained with UF method lead to acceptable coefficients of variation in comparison with standard method results (**Graph 6**). Coefficients of variation outside acceptable limits are only observed for low cell counts (percentage of mononuclear cells under 50%). There were 18 samples for which we obtained a coefficient of variation outside of acceptable limits: 7 AF, 5 PF, 2 PRF, 2 CSF, 1 SF and 1 PDF. Among them, 4 AF and 3 PF had abnormal cells on their smear. Both PRF samples had WBC counts above 100,000/μL, and both CSF had WBC counts under 100/μL.

**Graph 6.**
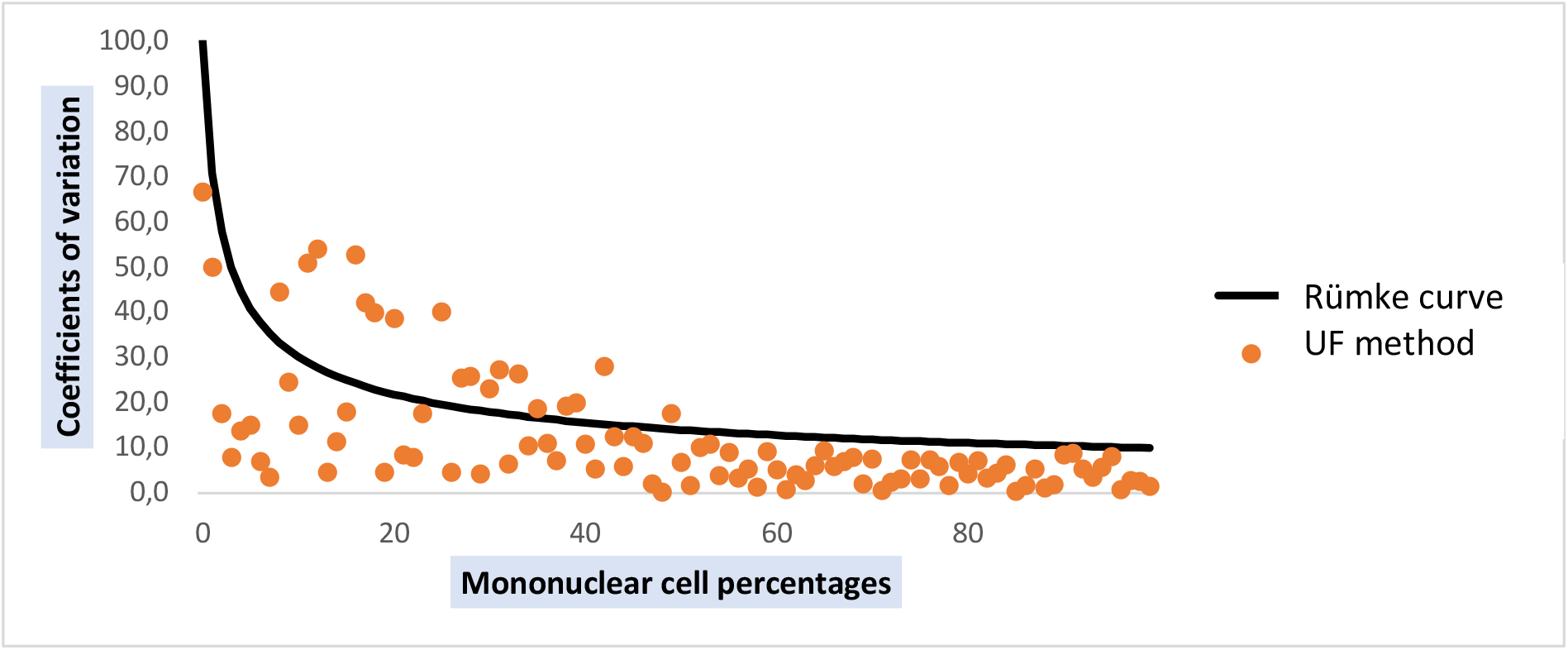
Comparison of mononuclear cell counts obtained by UF and standard methods. The Rümke curve was drawn using the Rümke table; it represents the limit value of acceptable coefficients of variation.

### 2/ Comparison of the performance of microbiological analysis methods

A total of 534 biological fluid samples were analysed by FCM (bacterial counts) using the SYSMEX UF4000 (Sysmex, Kobe, Japan). They were subsequently processed by routine microbiological procedures. Seven samples were excluded due to a lack of clinical or biological information and 1 sample which was only positive for *Candida spp*. was also excluded. Finally, 526 samples originating from 399 patients were included in this part of the study: 42 AF, 31 PF, 31 PRF, 16 PDF, 281 SF and 125 CSF. The mean age was 62 years (Interquartile Range (IQR): 24.5) and the sex ratio was 1.21. The characteristics of included patients are shown in **Table 5**.

**Table 5.**
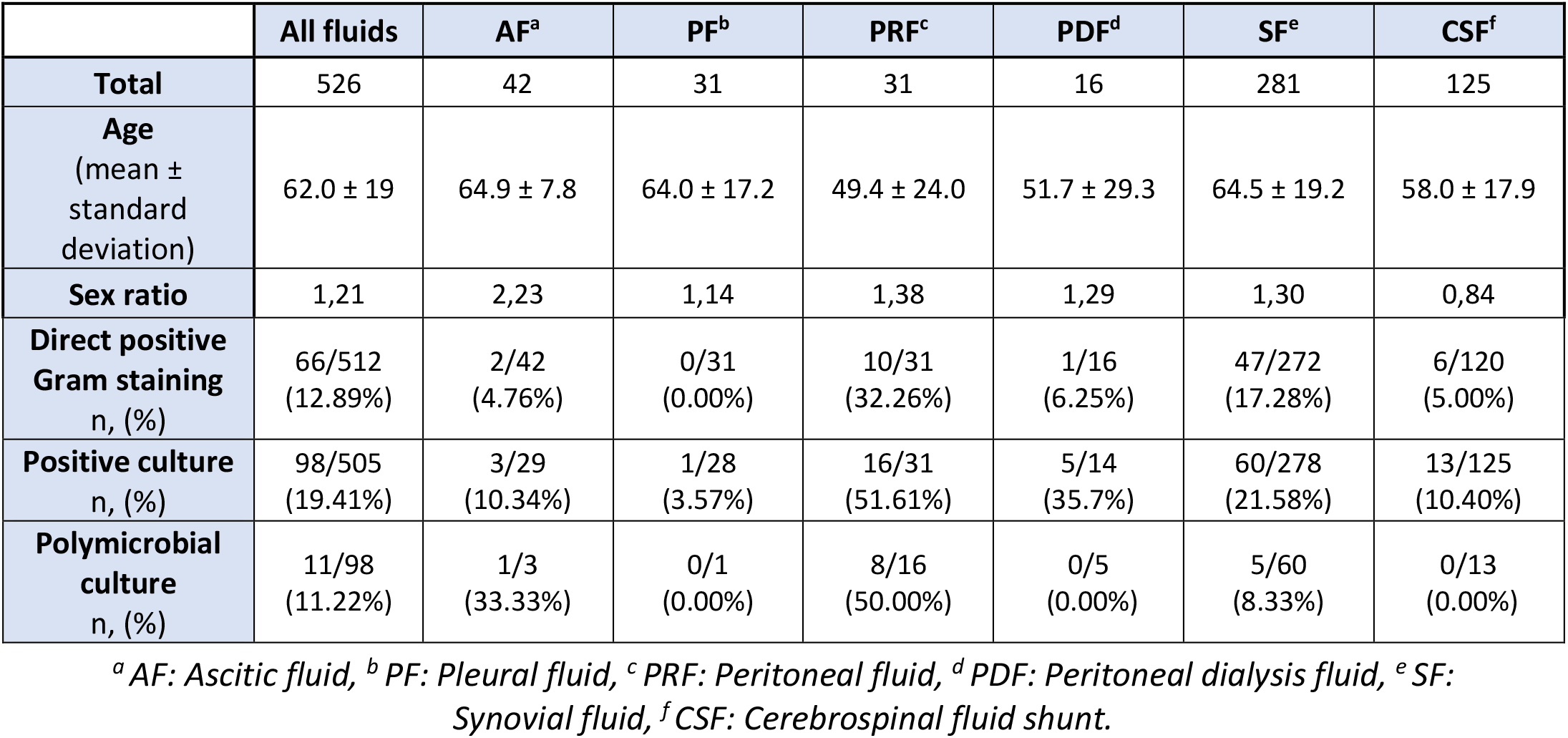
Characteristics of included patients for the comparison of the performance of microbiological analysis methods.

**Table 6** shows values of Bacteria and WBC counts estimated by the FCM of samples included according to culture positivity and DGS.

**Table 6.** Median and interquartile ranges of FCM parameters according to culture positivity and direct Gram staining.

| | Bacteria/μL | | | WBC/μL | | |
| --- | --- | --- | --- | --- | --- | --- |
|  | Median | IQR | p | Median | IQR | p |
| **Positive culture** | 382 | 64.65 – 2082.5 | p < 0.0001<sup>a</sup> | 3062.8 | 242.8 – 8757.75 | p < 0.0001<sup>a</sup> |
| **Negative culture** | 43 | 15 – 124.85 |  | 99.9 | 29.3 – 626.9 |  |
| **Direct positive Gram staining** | 635 | 190 – 3734.30 | p < 0.0001<sup>a</sup> | 3595 | 916.75 - 10096 | p < 0.0001<sup>a</sup> |
| **Direct negative Gram staining** | 43 | 15.8 – 126.25 |  | 100.4 | 29.5 - 651 |  |

<sup>a</sup> Mann-Whitney

DGS was performed in 512 samples, but culture was performed in only 505 samples (98.44%). DGS has a sensitivity of 50.00% and a specificity of 99.24% for the total of samples of our study (**Table 7**).

**Table 7.** Sensitivity and specificity of DGS according to culture results.

|  | All fluids | PRF<sup>a</sup> | SF<sup>b</sup> | CSF<sup>c</sup> |
| --- | --- | --- | --- | --- |
| **Sensitivity (%)** | 50.00 | 62.5 | 51.67 | 41.67 |
| **Specificity (%)** | 99.24 | 100 | 99.52 | 99.07 |

<sup>a</sup> PRF: peritoneal fluid, <sup>b</sup> SF: synovial fluid, <sup>c</sup> CSF: cerebrospinal fluid shunt

The ROC Curves of bacteria/μL and DGS positivity yielded an area under the curve (AUC) of 0.97, 0.88 and 0.89 for PRF, SF and CSF, respectively. The ROC curves for PDF, AF and PF were not calculated due to the low number of PDF included and because of the low number of positive samples among AF and PF (**Table 8**).

**Table 8.** Areas under the curve and optimal cut-off points for bacterial count by FCM versus DGS positivity and culture positivity.

|  |  | All fluids | PRF<sup>a</sup> | SF<sup>b</sup> | CSF<sup>c</sup> |
| --- | --- | --- | --- | --- | --- |
| **Direct Gram stain** | AUC | 0.88 | 0.97 | 0.88 | 0.89 |
|  | Optimal Cut-off point (bacteria/μL) | 185.0 | 465.0 | 1200.0 | 17.2 |
|  | Sensitivity / Specificity (%) | 77.8 / 82.5 | 100 / 85.7 | 95.7 / 67.1 | 100 / 54.4 |
|  | PPV / NPV (%) | 42.1 / 95.8 | 76.9 / 100 | 37.8 / 98.7 | 10.3 / 100 |
| **Microbial culture** | AUC | 0.83 | 0.83 | 0.80 | 0.60 |
|  | Optimal Cut-off point (bacteria/μL) | 465.0 | 541.7 | 6340.0 | 57.0 |
|  | Sensitivity / Specificity (%) | 58.2 / 96.3 | 62.5 / 93.33 | 45.0 / 93.6 | 46.2 / 83.0 |
|  | PPV / NPV (%) | 79.2 / 96.5 | 90.9 / 70.0 | 65.9 / 86.1 | 24.0 / 93.0 |

<sup>a</sup> PRF: Peritoneal fluid, <sup>b</sup> SF: Synovial fluid, <sup>c</sup> CSF: Cerebrospinal fluid shunt

Applying the calculated cut-off point (465.0 bacteria/μL) to the PRF samples, all PRF samples with DGS positive results were obtained (**Graph 7**). Also, 3 PRF samples with bacterial counts over the cut-off value by FCM (1035.4, 712.7 and 496, respectively) and DGS negative results were obtained. The first had a polymicrobial culture (*E. coli, P. mirabilis* and anaerobic bacteria), while the second had a WBC count of 54402/μL and was taken from a patient with acute appendicitis who was under antimicrobial treatment (3^rd^ generation Cephalosporin + Metronidazole) when the sample was obtained. The third had a WBC count of 7146.7/μL and was taken from a patient with peritonitis who was under antimicrobial treatment (3^rd^ generation Cephalosporin + Metronidazole) when the sample was obtained. For these 3 patients, a final diagnosis of infection was established based on clinical symptoms and the infectious aetiology was confirmed by other samples (blood cultures or other biological fluids).

**Graph 7.**
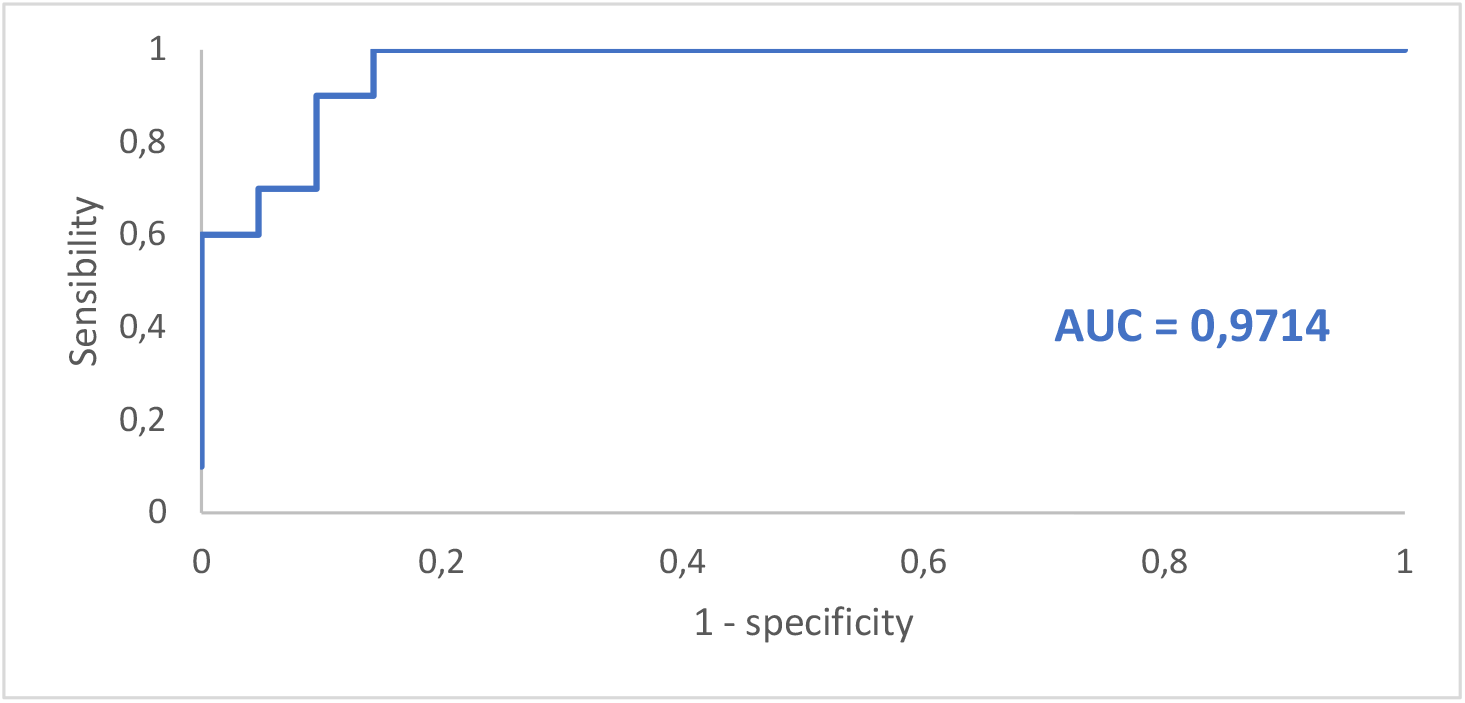
curve for bacterial counts by FCM versus DGS positivity for peritoneal fluid samples. AUC = Area under the curve.

Applying the calculated cut-off point (1200.0 bacteria/μL) to the SF samples, 45 of 47 SF samples with DGS-positive results were obtained (**Graph 8**). In the two patients with samples showing bacterial counts under the cut-off point and DGS positive results, a final diagnosis of infection was not established. Both had Gram positive cocci in DGS, but one had culture negative result and the other one had culture positive result (*S. capitis*). We suspected a false positive DGS result for the first and contamination of the microbial culture for the second. Seventy-four samples which showed high bacterial counts by FCM had DGS-negative results. Among these 74 samples, 9 were taken from patients with a final diagnosis of infection (established based on clinical symptoms). Moreover, two of them had culture positive results (*S. marcescens* and *S. lugduniensis*, respectively).

**Graph 8.**
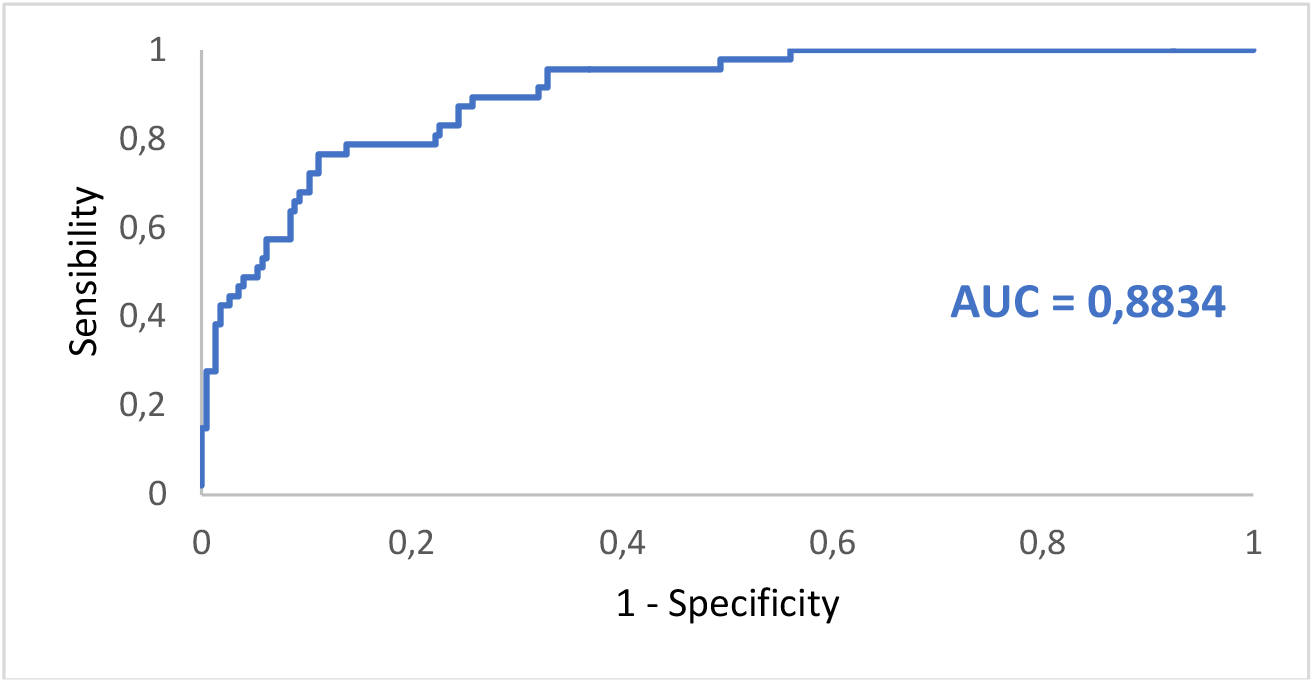
ROC curve for bacterial counts by FCM versus DGS positivity for synovial fluid samples. AUC = Area under the curve.

Applying the calculated cut-off point (17.2 bacteria/μL) to the CSF samples, all CSF samples with DGS-positive results were obtained (**Graph 9**). Fifty-two samples with high bacterial counts by FCM had DGS-negative results; six of them were taken from patients with a final diagnosis of infection (established based on clinical symptoms) and the infectious aetiology was confirmed on other samples (other CSF samples). Among these 6 samples, 3 had culture-positive results (*S. aureus, S. hominis* and *S. epidermidis*, respectively).

**Graph 9.**
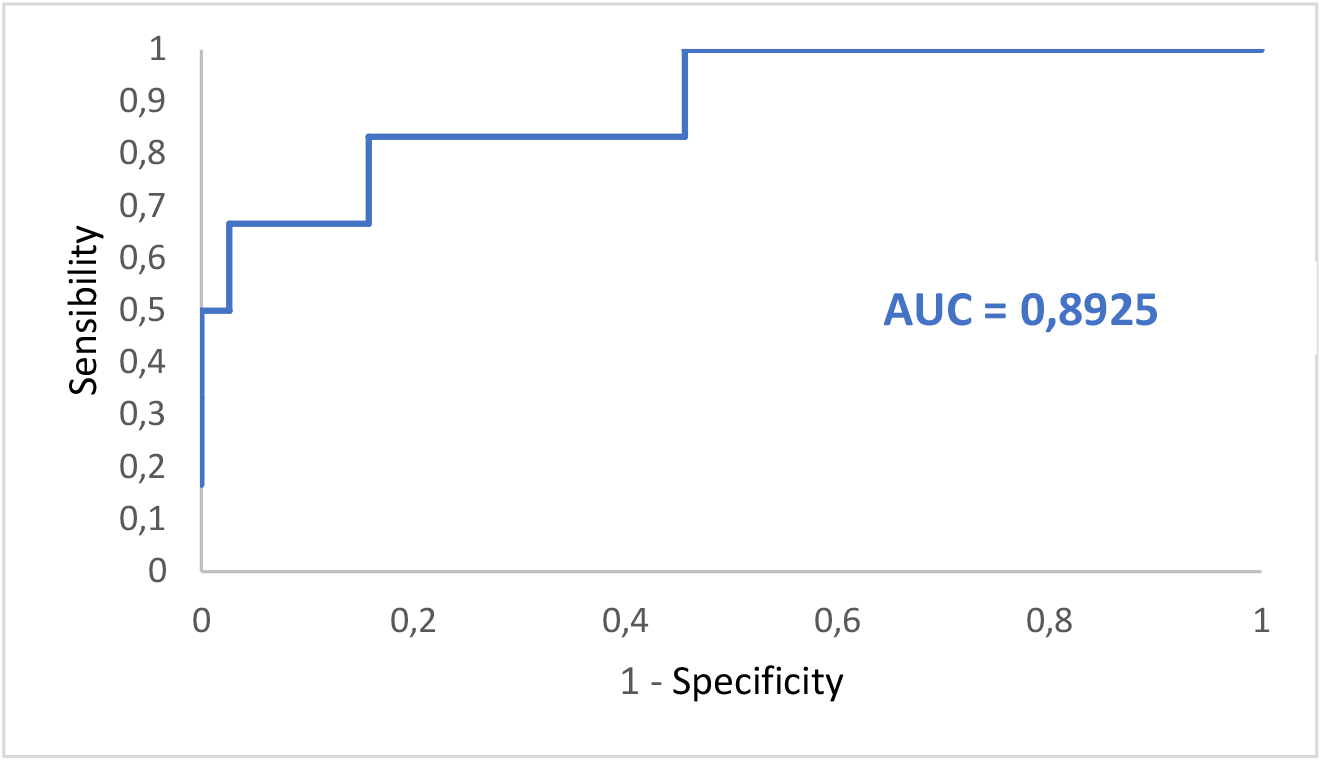
ROC curve for bacterial counts by FCM versus DGS positivity for cerebrospinal fluid samples. AUC = Area under the curve.

## Discussion

Cytobacteriological analysis of biological fluids is crucial in the management of patients suffering from effusions or infectious diseases. It requires cytological expertise for performing RBC counts, WBC counts, and differential leukocyte counts. It also requires microbiological expertise for performing DGS and for the interpretation of microbial cultures. We evaluated the flow cytometer UF4000 (Sysmex, Kobe, Japan) as a cytological and bacteriological analysis method in different biological fluid samples.

We compared the performance of cytological analyses of the UF4000 (Sysmex, Kobe, Japan) with the performance of the XN10 flow cytometer (Sysmex, Kobe, Japan) as well as that of standard reference methods (manual cell counts in KOVA counting chambers and manual differential leukocyte counts). With regard to cell counts, UF4000 (Sysmex, Kobe, Japan) showed a performance which was at least equivalent to those of the reference methods and superior to those of XN10 (Sysmex, Kobe, Japan). The Bland Altman plot of differences between RBC counts showed an average bias of −15.16/μL for the UF method. It is a weak bias; especially since there is no diagnostic threshold value for RBC counts. The appearance of the sample (haemorrhagic or not) is sufficient to allow a diagnostic orientation. An acceptable average bias was obtained for WBC counts (23.37/μL). Regarding the differential leukocyte counts, UF4000 (Sysmex, Kobe, Japan) showed excellent performance for body fluids which did not contain abnormal cells. Our results are in agreement with those of other studies, having demonstrated that the FCM allows to correctly classify and enumerate WBCs in the CSF, PRF and PDF^5–10^. Due to its characteristics, the XN10 (Sysmex, Kobe, Japan) is suitable for the analysis of samples containing a large quantity of RBC and WBC (> 10^3^ / μL). Therefore, the UF4000 (Sysmex, Kobe, Japan) seems more suitable for the cytological analysis of biological fluids which most often have low cell counts. However, it does not allow the differentiation of different types of leukocytes (lymphocytes, monocytes, polymorphonuclear neutrophils, eosinophils, polymorphonuclear basophils) and the detection of abnormal cells, which always makes it essential to carry out a manual smear for PF and AF. The main limitation of our study is the quantity of fluid sample necessary to perform the FCM analysis. Samples with a volume of less than 1 mL were systematically excluded (data not recorded), which limited our capacity for inclusion. Also, the standard method and flow cytometers had different limits of quantification and linearity ranges, which limited our ability to compare performance for body fluid samples with RBC or WBC counts with extreme values. Indeed, the automated methods had high limits of quantification which were markedly higher than those of the reference method.

Microbiological analysis results in biological fluids are tools for the management of patients suffering from infectious diseases. These results are crucial for the administration of targeted antimicrobial treatment and management of the infection. Requiring an incubation period, the results of microbial cultures can often only be obtained at least 24 hours after taking the sample; hence, the need for rapid detection methods for bacteria in biological fluid samples. We evaluated the performance of the flux cytometer UF4000 (Sysmex, Kobe, Japan) as a method for the rapid detection of bacteria in a variety of biological fluid samples and compared it with the culture and DGS results. There was a consensus between the bacterial count obtained by FCM and, DGS and culture results. Biological fluid samples with DGS-positive results had significantly higher bacterial and WBC counts than samples with DGS-negative results. Also, samples with culture-positive results had significantly higher bacterial and WBC counts than samples with culture-negative results. Our results are in agreement with previous studies which have demonstrated that bacterial count by FCM showed a good correlation with culture results^6,8,10,11^. We established optimal cut-off points for bacterial count by FCM in order to obtain DGS and culture-positive results. Several studies have suggested replacing the microbial culture with FCM for urine samples^12–14^. According to our results, biological fluid samples with low bacterial counts had a low probability of being culture-positive (NPV = 96.53%). However, due to the number of false-negative samples, the kind of patients involved, the precious nature of the samples and the potential severity of infections, the cessation of culture could not be considered in this type of sample. However, in order to determine the potential interest of FCM to replace DGS, we chose to determine specific and optimal cut-off points for each type of biological fluid sample so that they are as appropriate as possible. We have established 3 cut-off points for PRF (465.0 bacteria/μL), SF (1200.0 bacteria/μL) and CSF (17.2 bacteria/μL) with maximum sensitivity and negative predictive values. Unlike the DGS, the UF4000 flow cytometer (Sysmex, Kobe, Japan) does not provide an indication of the type of bacteria detected (Gram+, Gram-, cocci, bacilli). FCM is an automated and reliable method that could be used upstream of routine microbiological procedures. DGS may not be performed on biological fluid samples with bacterial counts below the defined cut-off points. In our study, this would have represented a total of 233 (47.3%) samples [18 PRF (58.1%), 153 SF (55.0%) and 62 CSF (51.7%)] for which the FCM could have replaced the DGS. This suggests that automating the bacteriological analysis of biological fluid samples could save significant technical time without losing efficiency. This work must be continued by other studies which will include greater numbers of fluid samples (especially PF, AF and PDF) in order to be able to determine optimal and specific cut-off points. In addition, the performance of mycological analyses of the UF4000 (Sysmex, Kobe, Japan) should be evaluated. Among the samples analysed, only 1 had DGS and culture-positive results with *C. albicans*; it was excluded from the study. This was an SF sample with WBC counts of 101,920/μL and “bacterial” counts above 1,000,000/μL. Therefore, it would be interesting to continue this work in order to evaluate the performance of the UF4000 (Sysmex, Kobe, Japan) as a method for the detection of yeasts in biological fluid samples. It has been shown that yeasts were sometimes confused with RBC^15^; hence, there is a need to re-analyse some samples on an additional module (UD10) to check for the presence of yeasts.

To the best of our knowledge, this is the first study to evaluate the performance of the UF4000 (Sysmex, Kobe, Japan) for the detection of bacteria in different types of biological fluid samples.

These results are promising for the organisation of microbiology laboratories and sampling circuits. The possibility of carrying out cytological and microbiological analyses of biological fluid samples on the same automated machine would simplify the sampling circuit (addressing the sample in a single laboratory, 24/7). It would also minimise the quantity of sample required (1mL for all of these analyses). However, the technical team will retain added value as cytological expertise for the search of abnormal cells or microbiological expertise for characterisation of the type of bacteria detected. A “flag” when detecting abnormal cells on the UF4000 (Sysmex, Kobe, Japan) is being investigated by the supplier.

In conclusion, the bacterial count by FCM according to the UF4000 (Sysmex, Kobe, Japan) correlates with DGS and culture results. It could be used upstream of routine microbiological procedures to improve and accelerate the diagnosis of infection in biological fluid samples.

